# A drought stress-responsive metabolite malate modulates stomatal responses through G-protein-dependent pathway in grapevine and Arabidopsis

**DOI:** 10.1101/2024.04.02.587830

**Authors:** Yoshiharu Mimata, Ruhai Gong, Xuanxuan Pei, Guochen Qin, Wenxiu Ye

## Abstract

Drought stress is a significant environmental threat to global agricultural production and distribution. Plant adaptation to dehydration stress involves intricate biological processes with substantial changes in metabolite composition. In this study, we investigated the role of tricarboxylic acid (TCA) cycle metabolites in drought tolerance in grapevine and Arabidopsis by metabolome, live cell imaging, electrophysiological and pharmacological approaches. Metabolome analysis revealed that amount of malate, citrate, and isocitrate increased over time in detached grapevine leaves. Ca^2+^ imaging and ion channel measurements indicated that fumarate, malate, and α-ketoglutarate induced cytosolic free Ca^2+^ concentration ([Ca^2+^]_cyt_) elevation in guard cells and directly activated a guard-cell anion channel SLAC1. However, only malate induced stomatal closure, which required increases in [Ca^2+^]_cyt_ in guard cells and activation of SLAC1. Through pharmacological experiments and reverse genetics analyses, G-proteins were identified as essential components of malate signaling by regulating second messenger production. These results indicate that TCA cycle metabolites are sensed individually by guard cells and that malate plays a key role in connecting metabolic regulation and drought tolerance through G-protein-dependent signal cascades.

## Introduction

Grapevine (*Vitis*) is one of the oldest domesticated crops and holds crucial economic importance for industries through the production of wine, brandy, juice, table grapes, and raisins. Despite the increasing demand for grapes and grape products, the global vineyard area is diminishing annually. In 2023, wine production was anticipated to reach its lowest levels in 60 years, primarily due to the impacts of global climate change (http://www.oiv.int/). Drought constitutes a major environmental stress with global implications for crop survival and yields. The regulation of metabolism stands out as a key mechanism for maintaining cell osmotic potential during drought stress. The metabolic responses to dehydration stress has been comprehensively studied in *Arabidopsis thaliana*. The metabolic reprogramming triggered by drought leads to elevated tricarboxylic acid (TCA) cycle intermediates in leaves (Urano et al., 2009; Pires et al., 2016). TCA cycle intermediates play an essential role in providing energy fuels and metabolic precursors. Malate is crucial due to its significant associations with stomatal movements, aluminum toxicity, CO_2_ fixation, ripening, and the taste of berries. Particularly in grapevines, malate is instrumental in determining wine quality and facilitating the growth of microorganisms for vinification (Fernie and Martinoia, 2009; Sweetman et al., 2009). The accumulation of malate is induced by environmental changes and may be linked to physiological responses in various tissues such as leaves, xylem, roots, and mesocarp (Van Kirk and Raschke, 1978; Kondo and Murata, 1987; Delhaize et al., 1993; Hedrich et al., 1994; Patonnier, 1999; Wada et al., 2008; Malcheska et al., 2017). However, the regulatory mechanism of physiological responses by TCA cycle metabolites remains unclear.

In response to drought, plants synthesize a phytohormone abscisic acid (ABA) and close stomatal pores, formed by pairs of guard cells in the epidermis of leaves, to prevent excessive water loss through guard cell signaling (Hetherington and Woodward, 2003; Murata et al., 2015). Stomatal closure is initiated by the transport of anions across the plasma membrane of guard cells through the slow-type anion channel encoded by the *SLOW ANION CHANNEL-ASSOCIATED 1* (*SLAC1*) gene (Schroeder et al., 2001; Negi et al., 2008; Vahisalu et al., 2008). In ABA signaling, SLAC1 is phosphorylated and activated by cytosolic Ca^2+^ sensor kinases, CALCIUM-DEPENDENT PROTEIN KINASEs (CDPKs) (Brandt et al., 2015), and a Ca^2+^-independent protein kinase OPEN STOMATA1 (OST1) (Geiger et al., 2009), leading to a decrease in turgor pressure and subsequent stomatal closure. Cytosolic free Ca^2+^ acts as a ubiquitous second messenger, and its concentration transiently increases in response to environmental, developmental, and growth signals (Luan and Wang, 2021). The increase in cytosolic free Ca^2+^ concentration ([Ca^2+^]_cyt_) results from the uptake of Ca^2+^ into the cell and the release of Ca^2+^ from internal stores through Ca^2+^ channels in response to a membrane potential shift (Hamilton et al., 2000) and second messengers such as cyclic adenosine diphosphate ribose (cADPR) (Leckie et al., 1998), cyclic adenosine monophosphate (cAMP) (Lemtiri-Chlieh and Berkowitz, 2004), inositol trisphosphate (IP_3_) (Gilroy et al., 1990), reactive oxygen species (ROS) (Pei et al., 2000), nitric oxide (NO) (Garcia-Mata et al., 2003), cyclic guanosine monophosphate (cGMP) (Wang et al., 2013), and nicotinic acid adenine dinucleotide phosphate (NAADP) (Navazio et al., 2000). In guard cells, cytosolic Ca^2+^ binds to the EF hands of CDPKs, leading to activation of CDPKs, which then induces stomatal closure through the phosphorylation of SLAC1 (Brandt et al., 2015).

Heterotrimeric G-proteins, composed of Gα, Gβ, and Gγ subunits, play pivotal roles in the generation of second messengers, such as cADPR, cAMP, IP_3_, and ROS (Zhang et al., 2011; Jin et al., 2013), thereby participating in various biological processes such as growth, development, and responses to environmental stimuli (Jin et al., 2013; Pandey, 2020). The genome of *A. thaliana* encodes one canonical Gα (GPA1), one Gβ (AGB1), and three Gγ subunits (AGG1–AGG3). GPA1 and/or AGB1 are involved in activating Ca^2+^ channels, slow-type anion channels and K^+^ channels and regulating stomatal movements by controlling the production of second messengers in ABA and Ca^2+^ signaling of guard cells (Wang et al., 2001; Fan et al., 2008; Zhang et al., 2011; Jeon et al., 2019). Recent findings have demonstrated that TCA cycle intermediates modulate systemic energy metabolism as metabolic signals “metabokine” *via* G-protein signaling cascades (Krzak et al., 2021). Malate as well as succinate directly binds to a G-protein-coupled receptor (GPCR) and causes rapid increases in [Ca^2+^]_cyt_ and IP_3_ accumulation (Trauelsen et al., 2017). However, it is unknown whether TCA cycle metabolites regulate G-protein signaling in plants.

In this study, we demonstrate that several TCA cycle metabolites accumulate in grapevine leaves during dehydration stress, among which malate most effectively regulates stomatal response *via* a G-protein signaling cascade. We propose that malate forms a hub between energy homeostasis and stress response.

## Results

### Metabolic responses of TCA cycle metabolites to dehydration stress in grapevine leaves

To investigate metabolic changes in response to drought, grapevine (*Vitis vinifera*) leaves were subjected to water-deficit stress. Detached leaves were sampled at 0–24 h of the dehydration stress treatment and subjected to non-targeted metabolome analysis. Principal component analysis (PCA) revealed that metabolite level changes were not pronounced between 0 and 1 h but gradually increased thereafter (Fig. **1A**). Significant metabolite changes were classified into three patterns: gradual decrease (subclass 1), gradual increase (subclass 2), and increase followed by decrease (subclass 3) (Supplementary Fig. **S1A**). The endogenous level of a phytohormone abscisic acid (ABA), an indicator of drought stress, increased after 2 h of dehydration stress and reached a plateau after 6 h (Fig. **1B**). After 24 h of treatment, out of 2,407 metabolites, 436 were upregulated, and 80 were downregulated (Fig. **1C**, Supplementary Table **S1**). Amino acids, lipids, terpenoids, phenolic acids, alkaloids, and flavonoids were the primary categories exhibiting changes (Fig. **1D**, Supplementary Fig. **S1B**). Among the upregulated metabolites, we specifically examined three TCA cycle metabolites—malate, isocitrate, and citrate—since malate functions as a signaling molecule inducing stomatal closure in *A. thaliana* (Mimata et al., 2022b). Malate, isocitrate, and citrate increased after 12 or 24 h of dehydration stress (Fig. **1E**). In contrast, *cis*-aconitate decreased immediately after 2–4 h and then returned to basal values.

**Figure 1.**
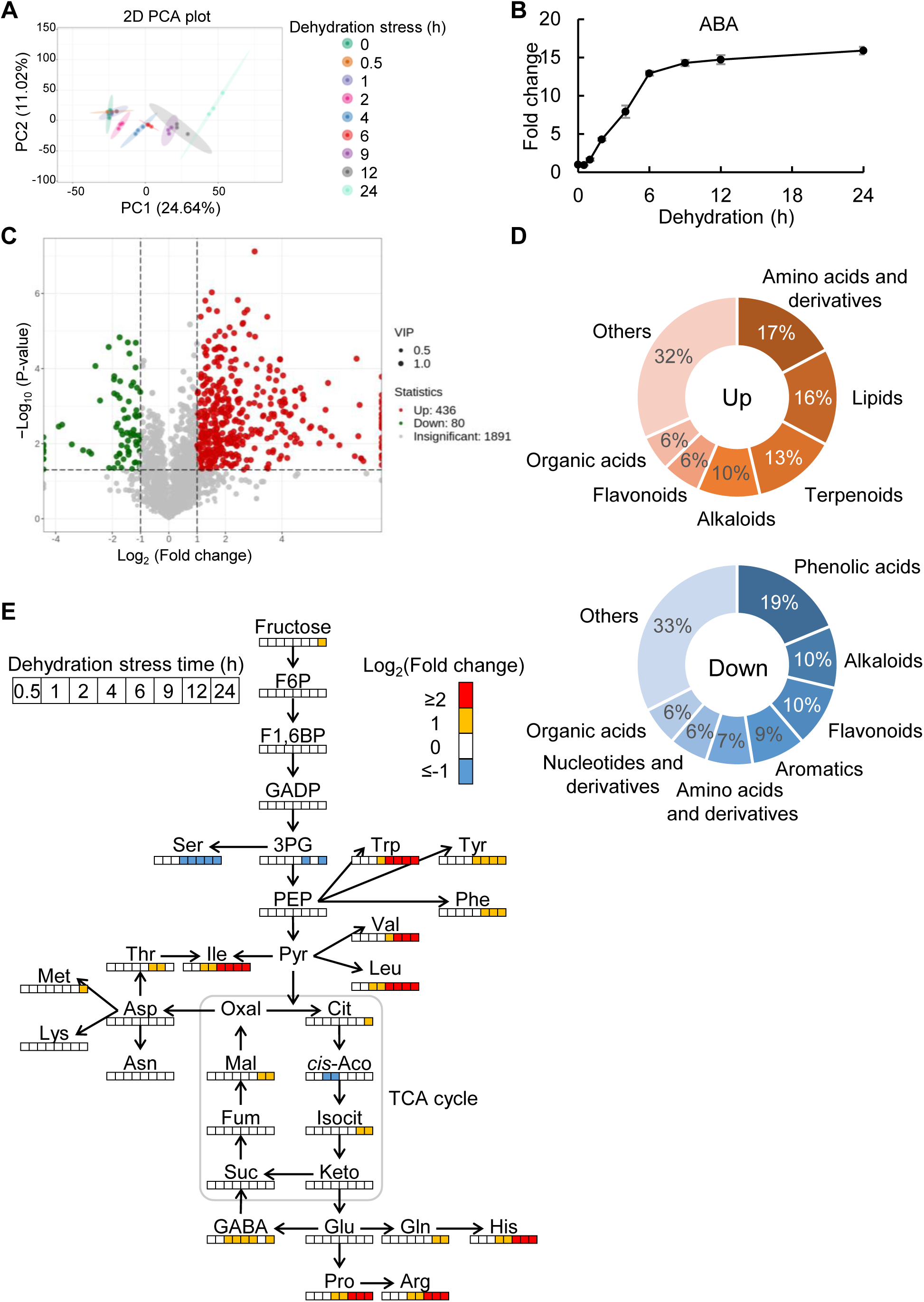
Metabolome analysis in grapevine leaves during dehydration treatment. A) PCA score plot of metabolomic datasets colored by the time of dehydration stress as clusters. Dots represent biological replicates. Ellipse display 95% confidence regions of each cluster. B) Relative ABA levels. Fold changes were normalized to the values of 0 h. Data are the mean ± SE. C) Volcano plot for the differential metabolites. Red and green dots mark the metabolites with significantly increased and decreased level in 0 h versus 24 h, respectively. Upregulated metabolites are defined VIP >1, log_2_(Fold change) ≥1 and P-value >0.05, downregulated metabolites are defined VIP >1, log_2_(Fold change) ≤−1 and P-value >0.05. P-value was calculated by Welch’s t-test. D) Categorization of the differential metabolites in **C)**. Upper panel shows upregulated metabolites, and lower panel shows downregulated metabolites. E) Relative TCA cycle metabolite levels at different time points. Heat maps represent log_2_(Fold change). Fold changes were normalized to the values of 0 h. Data were obtained from three independent biological replicates. Abbreviations: F6P, fructose-6-phosphate; F1,6BP, fructose 1,6-bisphosphate; GADP, glyceraldehyde 3-phosphate; 3PG, 3-phosphoglycerate; PEP, phosphoenolpyruvate; Pyr, pyruvate; Cit, citrate; *cis*-Aco, *cis*-aconitate; Isocit, isocitrate; Keto, α-ketoglutarate; Suc, succinate; Fum, fumarate; Mal, malate; Oxal, oxalacetate; Ser, serine; Trp, tryptophan; Tyr, tyrosine; Phe, phenylalanine; Val, valine; Leu, leucine; Ile, isoleucine; Thr, threonine; Asp, aspartate; Asn, asparagine; Met, methionine; Lys, lysine; Glu, glutamate; Gln, glutamine; His, histidine; Pro, proline; Arg, arginine; GABA, γ-aminobutyrate.

### Effects of TCA cycle metabolites on the [Ca^2+^]_cyt_ in guard cells

Since Ca^2+^ is a critical second messenger in guard cell signaling, we investigated the effects of TCA cycle metabolites (succinate, fumarate, malate, oxalacetate, α-ketoglutarate, citrate, *cis*-aconitate and isocitrate) and their associated compounds (acetate and pyruvate) on the elevations of [Ca^2+^]_cyt_ through live cell imaging of *A. thaliana* guard cells expressing a Ca^2+^ sensor fluorescent protein Yellow Cameleon 3.6. Exogenous application of fumarate, malate, α-ketoglutarate, acetate, and pyruvate induced elevations in [Ca^2+^]_cyt_, significantly increasing the frequency of [Ca^2+^]_cyt_ elevation compared to mock (Fig. **2**). This finding suggests that α-hydroxy or α-keto acids are effective in [Ca^2+^]_cyt_ elevation. Here, acetate caused transient long-term [Ca^2+^]_cyt_ increase (Fig. **2A**). Unlike the normal function of YC3.6, the CFP fluorescence did not return to its baseline, and the YFP fluorescence dropped below the basal level (Supplementary Fig. **S2)**. This dysfunctional response of YC3.6 following the Ca^2+^ surge served as an indicator of impending cell death (Ye et al., 2020). The Ca^2+^ channel blocker La^3+^ completely suppressed the malate-induced [Ca^2+^]_cyt_ increases (Fig. **2A, D**), indicating that Ca^2+^ channels are responsible for this Ca^2+^ response.

**Figure 2.**
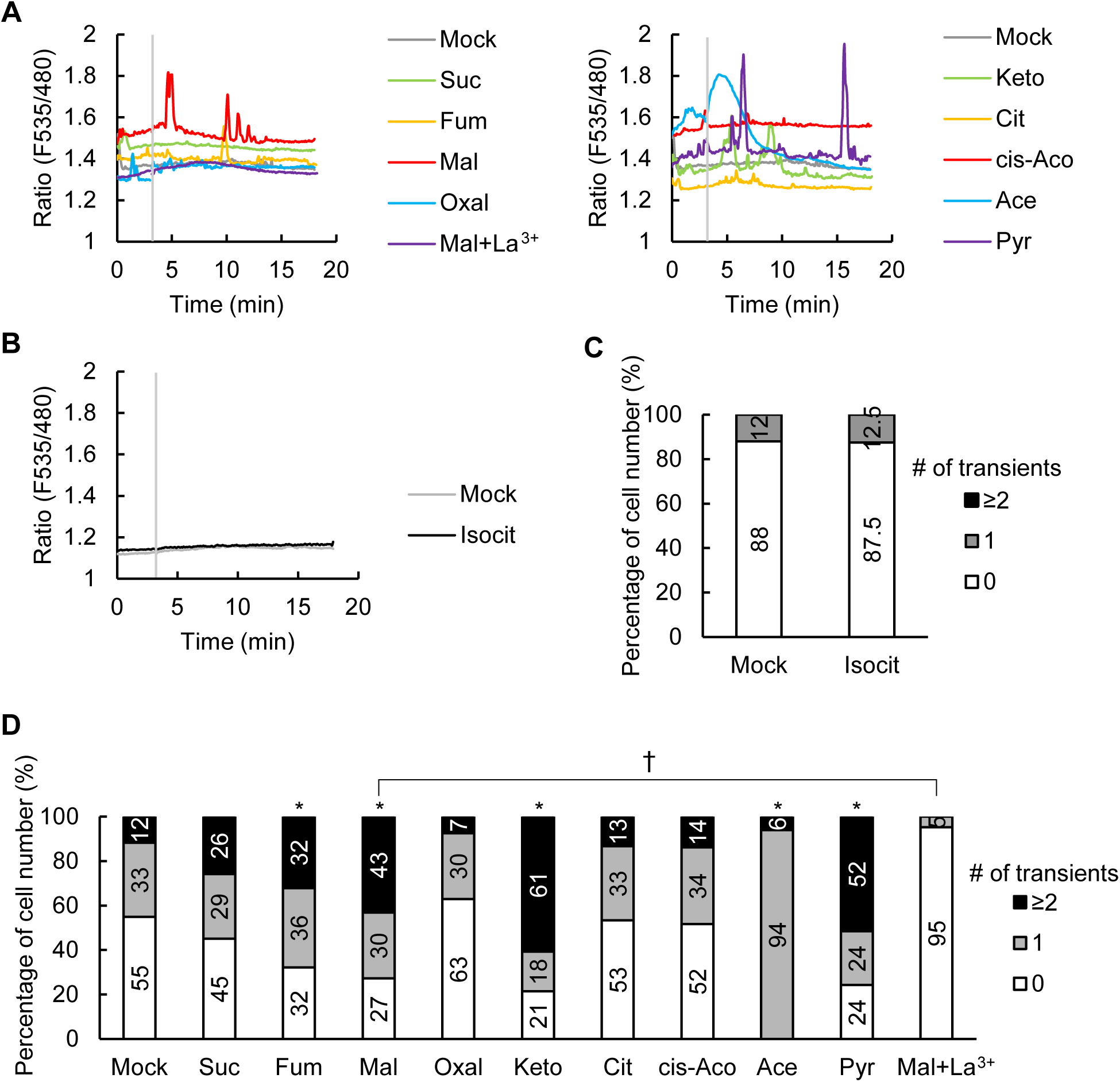
Ca^2+^ response to TCA cycle metabolites in guard cells. **A** and **B)** Representative traces of fluorescence emission ratios (535/480 nm) in *A. thaliana* guard cells expressing the Ca^2+^ sensor Yellow Cameleon 3.6. Grey bars indicate the time point when treatment was applied. The guard cells were treated with TCA cycle metabolites 3 min after the measurement. **C** and **D)** Percentage of number of guard cells showing different numbers of transient [Ca^2+^]_cyt_ increases. An increase in [Ca^2+^]_cyt_ is defined by an increase in fluorescence ratio by ≥0.1 U from the baseline ratio. Data were obtained from Mock (25), Isocit (16) for **C)**; Mock (51 guard cells), Suc (31), Fum (28), Mal (44) Oxal (27), Keto (28), Cit (30), cis-Aco (29), Ace (33), Pyr (33), Mal+La^3+^ (21) for **D)**. Asterisks and taggers indicate statistical significances based on Fisher’s exact test, P ≤0.05. Abbreviations: Isocit, isocitrate; Suc, succinate; Fum, fumarate; Mal, malate; Oxal, oxalacetate; Keto, α-ketoglutarate; Cit, citrate; cis-Aco, *cis*-aconitate; Ace, acetate; Pyr, pyruvate.

### Effects of TCA cycle metabolites on the activation of SLAC1 expressed in Xenopus oocytes

The activation of the SLAC1 anion channel plays a critical role in stomatal closure. VvSLAC1 and AtSLAC1 share 71% amino acid identity. A phenylalanine residue essential for pore gating (F450 in AtSLAC1) is conserved as F440 in VvSLAC1 (Qin et al., 2024). We predicted the structure of VvSLAC1 using *in silico* modeling. The modeling results showed that VvSLAC1 has ten transmembrane helices (Fig. **3** **A–D**). The pore is surrounded by an odd number of transmembrane helices and is occluded by F440. To further analyze VvSLAC1 activity, we conducted two-electrode voltage-clamp experiments on Xenopus oocytes. The negative currents in oocytes expressing VvSLAC1 were minimal, whereas those in oocytes expressing the VvSLAC1F440A mutant were much higher (Fig. **3E–G**). This indicates that VvSLAC1 is in an inactive state, while VvSLAC1F440A is constitutively active. This finding aligns with previous reports showing that an open-gate mutant, AtSLAC1F450A, exhibits substantial basal activity (Chen et al., 2010). The reversal potential was near the calculated equilibrium potential of Cl^−^ (approximately 50–80 mV), suggesting that VvSLAC1 permeates Cl^−^ and that these currents are minimally affected by leak currents.

**Figure 3.**
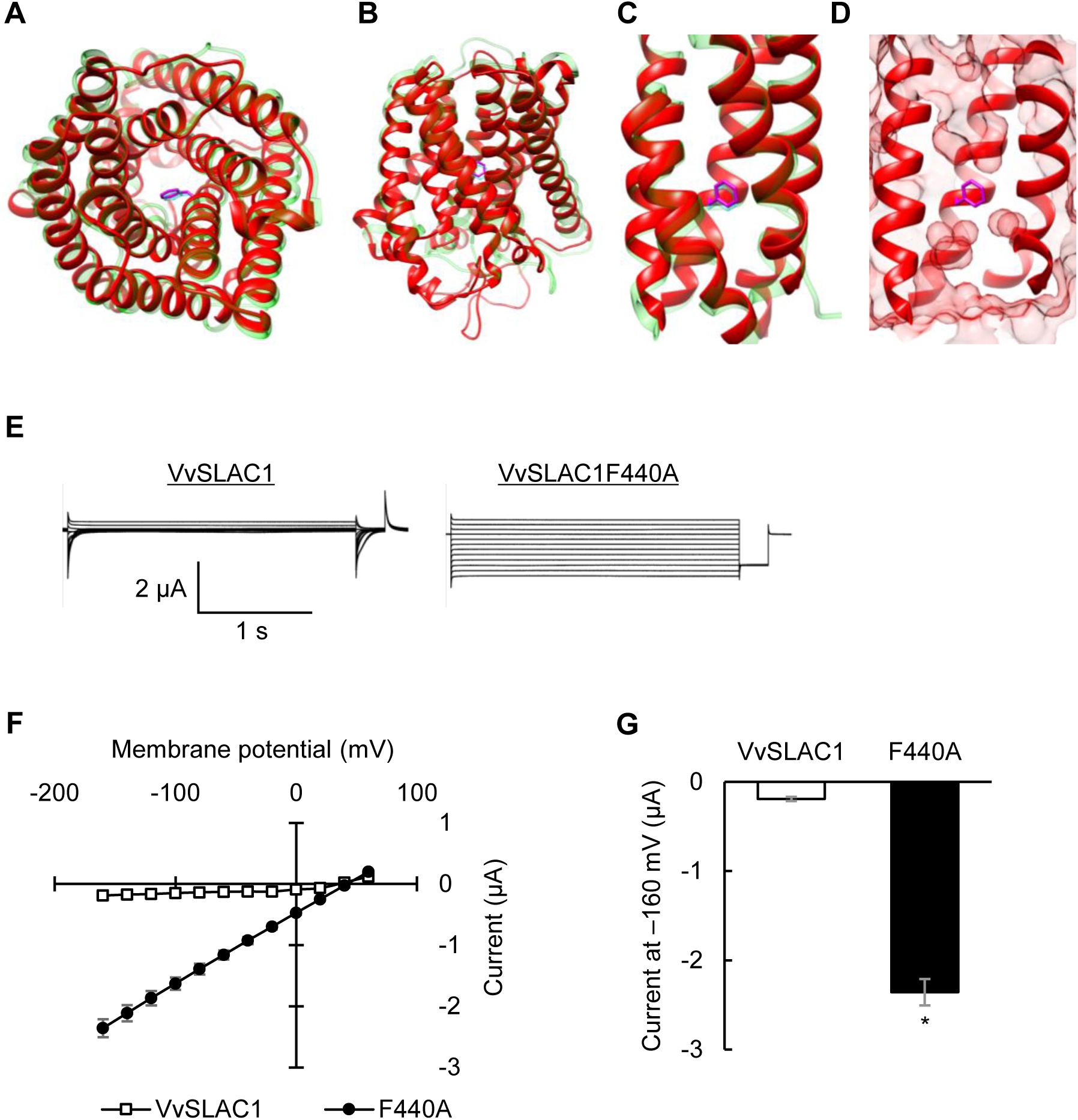
The negative currents of Xenopus oocytes expressing VvSLAC1F440A. **A to D)** A ribbon model of a VvSLAC1 protomer. A top view of the promoter is shown **A)**, and side views are shown **B to D)**. The VvSLAC1 structure model is displayed in red and the AtSLAC1 structure is displayed in green for comparison. The side chain of Phe 440 of VvSLAC1 is shown in magenta and Phe 450 of AtSLAC1 is shown in blue. The pore is shown as solid surface. **E)** Representative whole-cell negative current recordings in Xenopus oocytes expressing VvSLAC1F440A. The voltage pulse was commanded to clamp the membrane potential from +60 mV to −160 mV in −20 mV steps for 2.5 seconds with a holding potential of 0 mV. **F)** Average steady-state current–voltage curves of whole-cell negative current recordings. **G)** Average steady-state negative currents at −160 mV in **C)**. Data are the mean ± SE (n = 12 for VvSLAC1; n = 7 for VvSLAC1F440A). Different letters indicate statistical significances based on Student’s *t*-test, P < 0.05.

The activity of AtSLAC1F450A is enhanced by malate, whereas that of the wild-type AtSLAC1 is not (Mimata et al., 2022b). To examine the effects of TCA cycle metabolites, VvSLAC1 activity was continuously monitored during perfusion with a bathing solution supplemented with the metabolites. Isocitrate and citrate increased the negative currents in water-injected oocytes (Fig. **4A, B**), indicating that this activation is due to Xenopus endogenous transporters. None of the other tested metabolites affected the activity of wild-type VvSLAC1 (Fig. **4C, D**). Dicarboxylates, however, promoted the activity of VvSLAC1F440A. These results indicate that the open state of VvSLAC1 is a prerequisite for the promotion of its activity by TCA cycle metabolites. Next, we examined whether these effects depend on membrane potential. The increase in current magnitude was greater as the membrane potential became more negative (Fig. **4E, F**). Excluding oxalacetate, dicarboxylates significantly enhanced VvSLAC1 activity without affecting its reversal potential. These results show that dicarboxylates in the TCA cycle primarily promote Cl^−^ transport through SLAC1 once it is in the active state.

**Figure 4.**
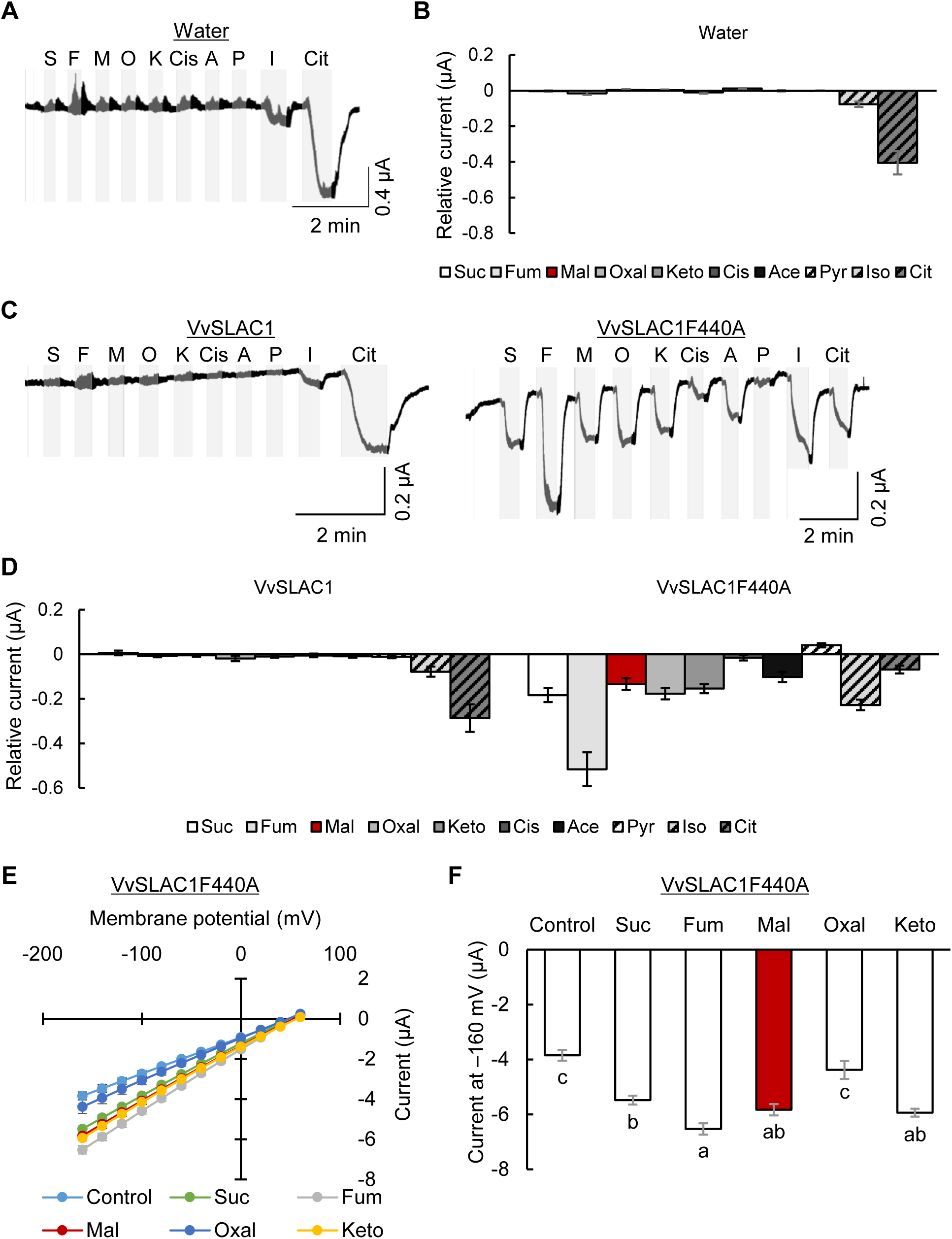
VvSLAC1 activity in the presence of TCA cycle metabolites. **A)** Representative whole-cell negative current traces during perfusion with TCA cycle metabolites in water-injected Xenopus oocytes. The voltage pulse was commanded to clamp the membrane potential at −120 mV. Grey regions indicate metabolite perfusion and white regions indicate washout. **B)** Average of relative currents. The currents were normalized to the mock-treated current. Data were obtained from six oocytes per condition. Data are the mean ± SE. **C)** Representative whole-cell negative current traces during perfusion with TCA cycle metabolites in Xenopus oocytes expressing VvSLAC1F440A. **D)** Average of relative currents. Data are the mean ± SE (n = 5 for VvSLAC1; n = 6 for VvSLAC1F440A). **E)** Average steady-state current–voltage curves of whole-cell negative current recordings in bathing solution supplemented with TCA cycle metabolites. The voltage pulse was commanded to clamp the membrane potential from +60 mV to −160 mV in −20 mV steps for 2.5 seconds with a holding potential of 0 mV. **F)** Average steady-state negative currents at −160 mV in **E)**. Data are the mean ± SE. Data were obtained from four oocytes per condition. Different letters indicate statistical significances based on one-way ANOVA with Tukey’s HSD test, P < 0.05. Abbreviations: Suc/S, succinate; Fum/F, fumarate; Mal/M, malate; Oxal/O, oxalacetate; Keto/K, α-ketoglutarate; Cit, citrate; Isocit/I, isocitrate; Cis, *cis*-aconitate; Ace/A, acetate; Pyr/P, pyruvate.

### Malate emerges as a specific modulator in the regulation of stomatal responses

To assess how TCA cycle metabolites influence plant responses to drought stress, we measured stomatal aperture in the presence of the metabolites. After stomata were fully open under light, each metabolite was applied, and stomatal aperture was measured. A significant reduction in stomatal aperture was observed exclusively with malate treatment in *V. vinifera* (Fig. **5A**). Consistent results were obtained using *A. thaliana* (Fig. **5B**). These findings suggest that malate specifically acts as a modulator of stomatal closure.

**Figure 5.**
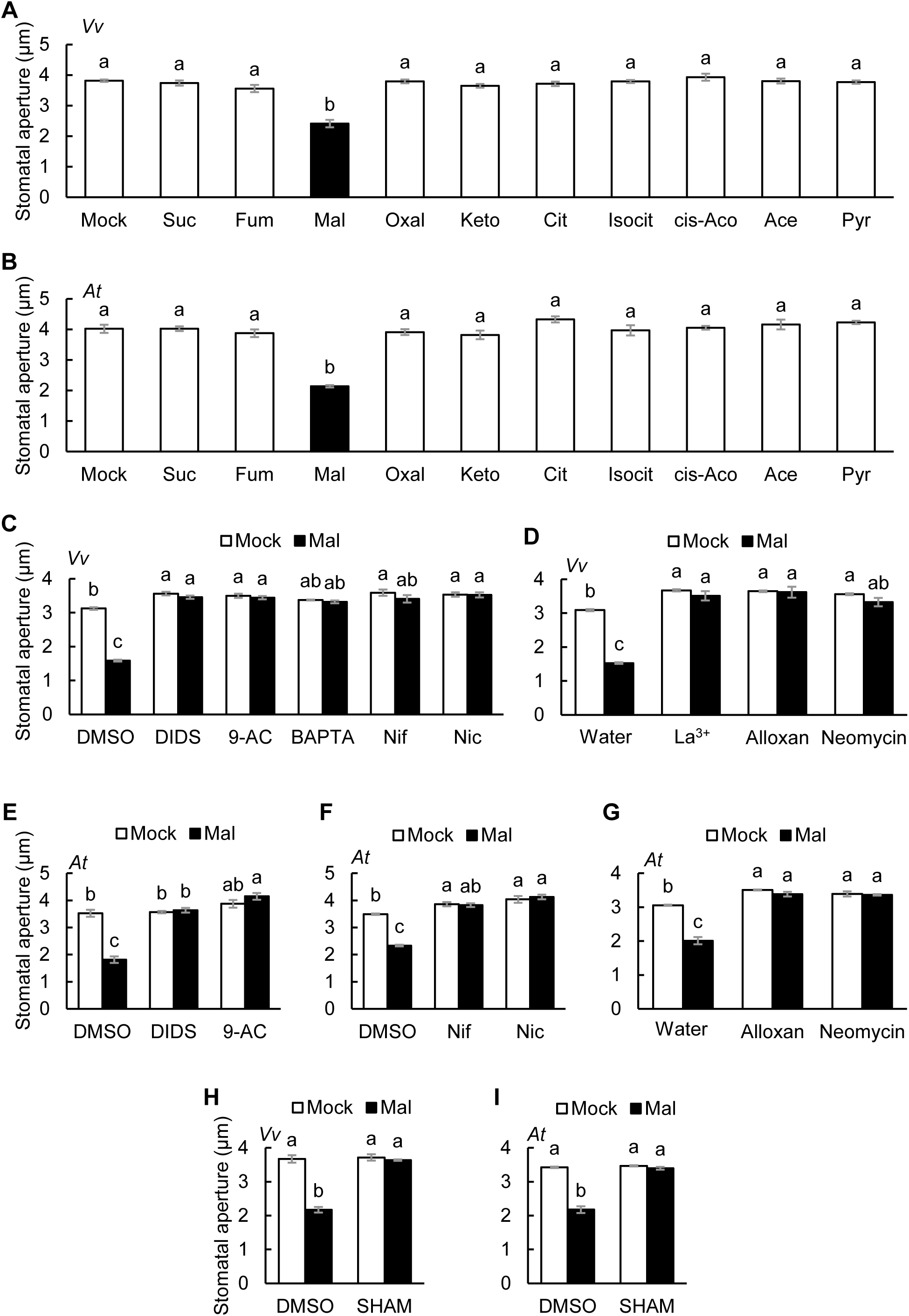
Malate-induced stomatal closure is mediated by anion channels and second messengers. **A** and **B)** Effects of TCA cycle metabolites on stomatal aperture in **A)** *V. vinifera* or **B)** *A. thaliana* leaves. Data are the mean ± SE. Different letters indicate statistical significances based on one-way ANOVA with Tukey’s HSD test, P <0.05. **C** to **I)** Effects of inhibitors on malate-induced stomatal closure in **C**, **D** and **H)** *V. vinifera* or **E** to **G**, **I)** *A. thaliana* leaves. Averages of stomatal apertures from four independent experiments (n = 4) are shown. Data are the mean ± SE. Different letters indicate statistical significances based on two-way ANOVA with Tukey’s HSD test, P < 0.05. Abbreviations: Suc, succinate; Fum, fumarate; Mal, malate; Oxal, oxalacetate; Keto, α-ketoglutarate; Cit, citrate; Isocit, isocitrate; cis-Aco, *cis*-aconitate; Ace, acetate; Pyr, pyruvate; Nif, nifedipine; Nic, nicotinamide.

To further investigate the malate signaling pathway, we conducted a series of pharmacological experiments (Supplementary Table **S2**). We applied anion channel blockers, DIDS and 9-anthracenecarboxylic acid (9-AC) (Schwartz et al., 1995; Geiger et al., 2009), extracellular Ca^2+^ chelator 1,2-bis(2-aminophenoxy)ethane-*N*,*N*,*N’*,*N’*-tetraacetic acid (BAPTA) (Levchenko et al., 2005), and Ca^2+^ channel blockers, nifedipine and La^3+^ (Reiss and Herth, 1985; Pei et al., 2000). Malate-induced stomatal closure in both *V. vinifera* and *A. thaliana* was abolished by all inhibitors (Fig. **5C–F**; Mimata *et al*., 2022b). These results suggest anion channels and Ca^2+^ signaling *via* Ca^2+^ channels are essential for malate-induced stomatal closure in *V. vinifera* and *A. thaliana*, consistent with the results for [Ca^2+^]_cyt_ and SLAC1 activity (Fig. **2A, D**, **4D–F**; Mimata *et al*., 2022b).

### Malate stimulates Ca^2+^ signaling *via* second messengers including cADPR, cAMP, and IP_3_

To further elucidate the malate-induced stomatal closure, the involvement of second messengers relevant to Ca^2+^ signaling, including cADPR, cAMP, IP_3_, ROS, NO, cGMP, NAADP, and PIP_3_, was investigated by a pharmacological approach. We applied inhibitors targeting these second messengers that are well-established in plant studies. Nicotinamide, alloxan, neomycin, and salicylhydroxamic acid (SHAM), which are inhibitors of cADPR (Dodd et al., 2007), cAMP (Ma et al., 2009), IP_3_ (Tang et al., 2007), and peroxidase-catalyzed ROS production (Mori et al., 2001), respectively, completely abolished the malate-induced stomatal closure in *V. vinifera* and *A. thaliana* (Fig. **4C**, **D**, **F****–I**). On the other hand, *N*-nitro-L-arginine methyl ester (L-NAME), 2-(4-carboxyphenyl)-4,4,5,5-tetramethylimidazoline-1-oxyl-3-oxide (cPTIO), LY83583, Ned 19, and wortmannin, which are an NO synthetase inhibitor (Joudoi et al., 2013), an NO scavenger (Isner et al., 2019), a guanylate cyclase inhibitor, an antagonist of NAADP (González et al., 2012), and an inhibitor of PIP_3_ production (Matsuoka et al., 1995), respectively, have little inhibitory effect on malate-induced stomatal closure (Supplementary Fig. **S3**). Ca^2+^ imaging experiments further showed that nicotinamide, alloxan, and neomycin but not SHAM inhibited malate-induced [Ca^2+^]_cyt_ oscillations in Arabidopsis (Fig. **6A, B**). These results suggest that Ca^2+^ signaling involving cADPR, cAMP, and IP_3_ is required for malate-induced stomatal closure.

**Figure 6.**
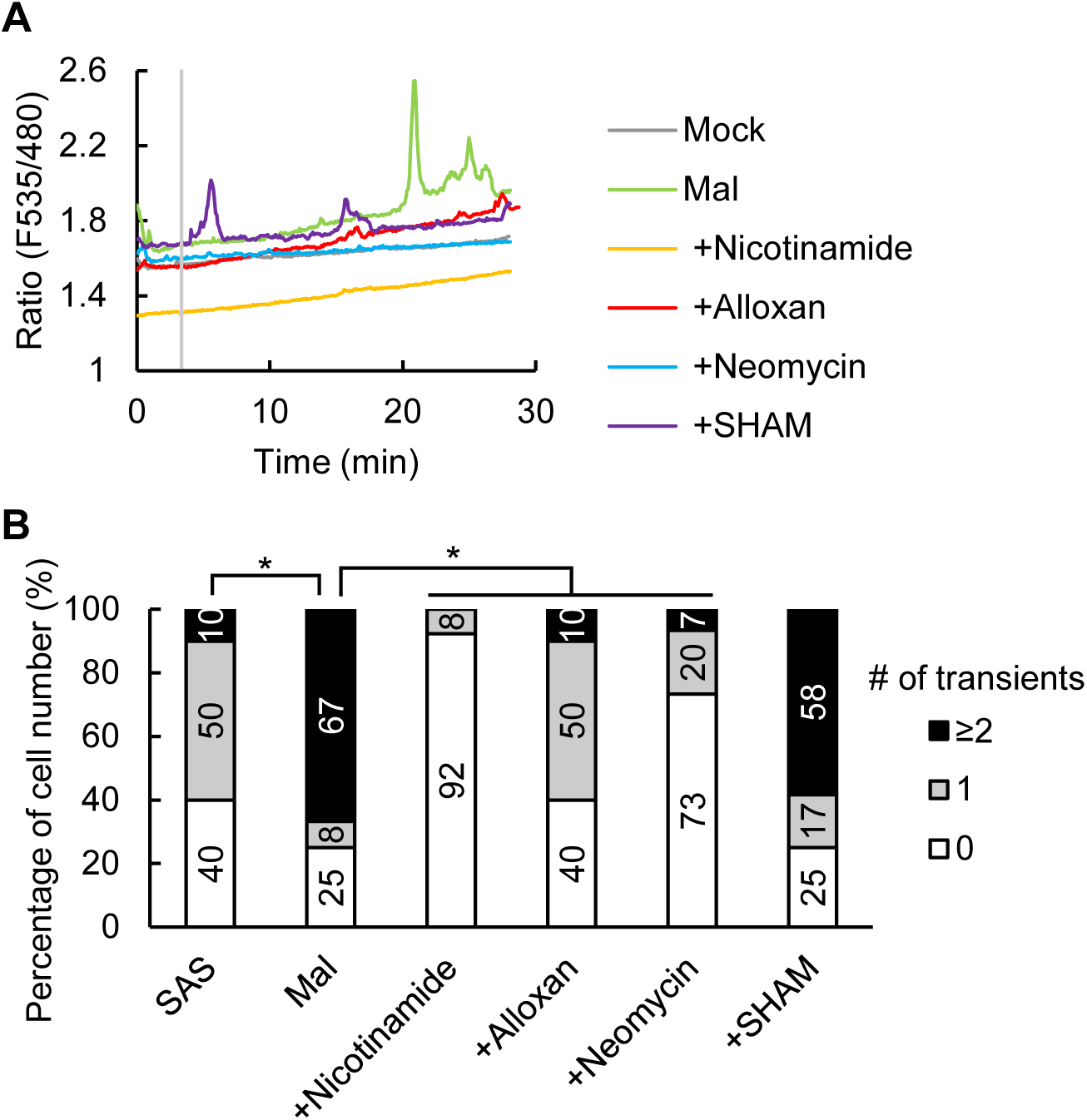
Malate-induced [Ca^2+^]_cyt_ elevations are mediated by cADPR, cAMP and IP_3_. A) Representative traces of fluorescence emission ratios (535/480 nm) in *A. thaliana* guard cells expressing the Ca^2+^ sensor Yellow Cameleon 3.6. Grey bar indicates the time point when treatment was applied. The guard cells were treated with malate 3 min after the measurement. Inhibitors were added 5 min before starting imaging. B) Percentage of number of guard cells showing different numbers of transient [Ca^2+^]_cyt_ increases. An increase in [Ca^2+^]_cyt_ is defined by an increase in fluorescence ratio by ≥0.1 U from the baseline ratio. Data were obtained from Mock (10 guard cells), Mal (12), +Nicotinamide (13), +Alloxan (10), +Neomycin (15), +SHAM (12). Asterisks indicate statistical significances based on Fisher’s exact test, P ≤0.05.

### G-proteins are master regulators in malate signaling

Since the generation of cADPR, cAMP, IP_3_, and ROS is regulated by G-proteins, their roles in the stomatal response to malate were investigated with the G-protein inhibitors. All inhibitors completely abolished the malate effect in *V. vinifera* and *A. thaliana* (Fig. **7A**, **B**, Supplementary Fig. **S3E, F**). To confirm the pharmacological result, a reverse genetic approach was performed using loss-of-function mutants for the Gα subunit and Gβ subunit, *gpa1* and *agb1*. The stomata of *gpa1* and *agb1* mutants were insensitive to malate (Fig. **7C**). GDPβS also suppressed malate-induced [Ca^2+^]_cyt_ elevations and ROS production (Fig. **7D–F**). Furthermore, ROS production was not promoted by malate in the *gpa1* and *agb1* mutants (Fig. **7G**). These data together demonstrate that malate signaling is transduced by G-proteins in guard cells.

**Figure 7.**
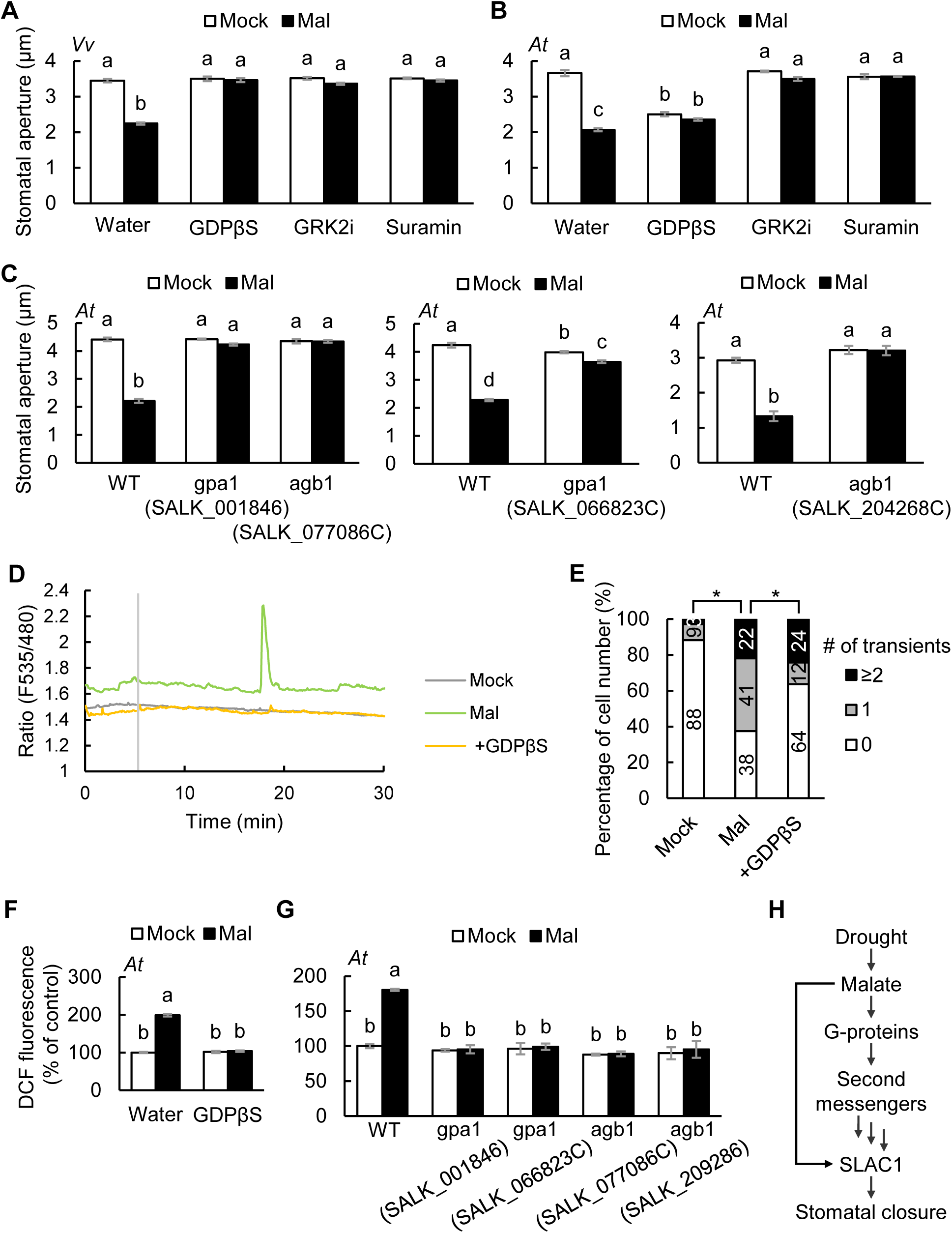
Malate signaling is mediated by G-proteins. **A and B)** Effects of G-protein inhibitors on malate-induced stomatal closure in **A)** *V. vinifera* or **B)** *A. thaliana* leaves. C) Effects of *gpa1* and *agb1* mutation on malate-induced stomatal closure. Averages of stomatal apertures from four independent experiments (n = 4) are shown. Data are the mean ± SE. Different letters indicate statistical significances based on two-way ANOVA with Tukey’s HSD test, P < 0.05. D) Representative traces of fluorescence emission ratios (535/480 nm) in *A. thaliana* guard cells expressing the Ca^2+^ sensor Yellow Cameleon 3.6. Grey bar indicates the time point when treatment was applied. The guard cells were treated with malate 5 min after the measurement. Inhibitors were added 5 min before starting imaging. E) Percentage of number of guard cells showing different numbers of transient [Ca^2+^]_cyt_ increases. An increase in [Ca^2+^]_cyt_ is defined by an increase in fluorescence ratio by ≥0.1 U from the baseline ratio. Data were obtained from Mock (34 guard cells), Mal (32), +GDPβS (33). Asterisks indicate statistical significances based on Fisher’s exact test, P ≤0.05. F) Effects of GDPβS on malate-induced ROS production in *A. thaliana* guard cells. The ROS-sensitive dye, 2’,7’-dichlorodihydrofluorescein diacetate (H_2_DCF-DA) was used for ROS detection in guard cells. Fluorescence intensity was normalized to mock value in water. G) Effects of *gpa1* and *agb1* mutation on malate-induced ROS production in *A. thaliana* guard cells. Fluorescence intensity was normalized to mock value in WT. Averages from three independent experiments (n = 3) are shown. Data are the mean ± SE. Different letters indicate statistical significances based on two-way ANOVA with Tukey’s HSD test, P <0.05. H) A proposed working model for malate signaling.

## Discussion

Drought has severe impacts on agricultural crops and results in metabolite fluctuations in plants. In this study, we identified 436 upregulated and 80 downregulated metabolites in grapevine leaves in response to dehydration stress (Fig. **1C**). These included modulators of stomatal movements, such as phytohormones, primary metabolites, and aromatic secondary metabolites (Supplementary Table **S1**). Among the upregulated metabolites, there were those that induce stomatal closure or inhibit stomatal opening, such as ABA, phaseic acid, adenosine-3’-5’-diphosphate, γ-aminobutyric acid (GABA), and malic acid. On the other hand, among the downregulated metabolites, there were those that promote stomatal opening or inhibit stomatal closure, such as indole-3-acetic acid and 5-aminolevulinic acid. Amino acids and TCA cycle metabolites are osmolytes and their accumulation reduces the water potential. Several amino acids induce stomatal closure through a pathway dependent on glutamate receptor-like channels (GLRs) (Kong et al., 2016). Although TCA cycle metabolites, especially malate, are key metabolites for stomatal movements, their role in signal transduction has remained largely unknown. This study specifically focused on the effect of TCA cycle metabolites and clarified the mechanism underlying stomatal closure triggered by TCA cycle metabolites.

### Specific guard cell responses triggered by TCA cycle metabolites

TCA cycle metabolites elicit distinct guard cell responses, including increases in [Ca^2+^]_cyt_, activation of SLAC1, and stomatal closure (Fig. **2, 4, 5**). The different specificity of the Ca^2+^ response to TCA cycle metabolites (Fig. **2**), compared to SLAC1 activation (Fig. **4**), suggests that SLAC1 activation does not necessarily lead to increases in [Ca^2+^]_cyt_, and a carboxylate receptor independent from SLAC1 may exist. Despite both α-ketoglutarate and oxaloacetate are α-keto acids, only α-ketoglutarate, with a carbon chain length similar to glutamate, was effective in inducing elevation of [Ca^2+^]_cyt_ (Fig. **2A, D**), indicating the importance of carbon chain length in perception. It was shown that dicarboxylate glutamate induces [Ca^2+^]_cyt_ elevation in guard cells and stomatal closure through a pathway dependent on glutamate receptor-like channels (GLRs), while malate signaling is independently of GLRs (Mimata et al., 2022b). It is plausible that α-ketoglutarate is recognized by GLRs. Even though fumarate and α-ketoglutarate caused Ca^2+^ elevation, SLAC1 activation, and ROS production (Fig. **2A**, **D**, **4**, Supplementary Fig. **S4A**), they failed to close the stomata (Fig. **5A, B**). An unknown factor specific to malate, but not to fumarate or α-ketoglutarate, likely contributes to this process and requires further investigation. Taken together, TCA cycle metabolites are individually sensed by guard cells through distinct mechanisms (Fig. **2, 4, 5**).

### Malate plays a key role in stress responses

Dehydration stress initiated the accumulation of ABA after 2 h (Fig. **1B**), which correlates with the onset of stomata closure (Hopper et al., 2014). The endogenous level of malate increased 12 h after dehydration stress (Fig. **1E**). Supporting this, a related study reported 6.78-fold increase in the relative abundance of malate in Arabidopsis aerial parts after 10 h of dehydration stress (Urano et al., 2009). These findings showed that malate slowly accumulates in whole leaves. Malate was observed to be secreted during stomatal closure (Van Kirk and Raschke, 1978), with its content in guard cells decreasing right after ABA treatment (Kondo and Murata, 1987; Jin et al., 2013). Consistent with these reports, the concentration of apoplast malate increased within 15 minutes in response to elevated CO_2_, which triggered stomatal closure (Hedrich et al., 1994). The apoplastic malate concentration is estimated to reach approximately 10 mM, with other metabolites being present in lower concentrations or undetected in leaves of several plant species (Lohaus et al., 1995; Gabriel and Kesselmeier, 1999; Hedrich et al., 2001). Notably, the malate exporter AtALMT12, mainly expressed in guard cells and localized at the plasma membrane, is activated through the ABA signaling pathway (Meyer et al., 2010; Sasaki et al., 2010), and loss-of-function mutation of AtALMT12 has been shown to increase malate content in leaves (Medeiros et al., 2016). These findings suggest that malate is rapidly expelled by transporters and gradually recharged, likely through intracellular biosynthesis, under stress conditions.

Malate was the most potent TCA cycle metabolite that induces stomatal closure (Fig. **5A, B**). The malate-induced stomatal closure has also been confirmed in other methods: feeding malate through the petiole decreases stomatal aperture and conductance in ash and aspen trees (Patonnier, 1999; Rasulov et al., 2018). The *atalmt12* mutants exhibit increased malate accumulation and weaker and slower stomatal closure in leaves (Meyer et al., 2010; Sasaki et al., 2010; Medeiros et al., 2016). As AtALMT12 is gated by malate (Meyer et al., 2010), exported malate accelerates malate efflux as a feedback loop. This process subsequently leads to malate accumulation in apoplast, which activates SLAC1 and drives stomatal closure. Moreover, exogenous application of malate inhibits stomatal opening (Esser et al., 1997). Therefore, malate may play a role in maintaining stomatal closure to reduce water loss and enhance drought tolerance. This hypothesis is further supported by a report showing that *atalmt12* mutants are sensitive to drought stress (Medeiros et al., 2016).

### Malate signaling is mediated by a specific set of second messengers

Stomatal measurements (Fig. **5D, G**) and Ca^2+^ imaging (Fig. **6**) with inhibitors suggest the involvement of second messengers including cAMP, cADPR, and IP_3_ in malate signaling. cAMP triggers Ca^2+^ influx through CYCLIC NUCLEOTIDE-GATED CHANNELS (CNGCs) in guard cells (Lemtiri-Chlieh and Berkowitz, 2004; Ali et al., 2007). Recently, it was reported that multiple CNGCs work redundantly as ABA-activated Ca^2+^ channels, which are necessary for ABA-induced Ca^2+^ oscillations and stomatal closure independently of ROS (Tan et al., 2023; Yang et al., 2024). Future studies should investigate the involvement of CNGCs in malate signaling.

Malate-induced stomatal closure required the peroxidase activity (Fig. **5H, I**). Malate promotes ROS production by peroxidases (Mimata et al., 2022a), and ROS increase [Ca^2+^]_cyt_ *via* plasma membrane Ca^2+^ channels (Pei et al., 2000). Based on these observations, we hypothesized that malate accelerates Ca^2+^ influx by promoting ROS production. Contrary to our hypothesis, the inhibition of ROS production did not impair malate-induced Ca^2+^ responses (Fig. **6**). Moreover, blocking Ca^2+^ influx did not affect ROS production (Supplementary Fig. **S4B**). These findings indicate that ROS production is independent of Ca^2+^ signaling in malate-induced stomatal closure.

### Malate signaling is transduced by G-protein signaling cascades

In mammals, carboxylates, such as succinate and malate, are sensed by a GPCR. Succinate is released from stimulated macrophages and injured tissues, reaching millimolar concentrations locally (Chouchani et al., 2014; Littlewood-Evans et al., 2016). Extracellular succinate activates GPCRs, stimulating IP_3_ formation, inhibiting cAMP production, and increasing [Ca^2+^]_cyt_ *via* the G-protein signaling pathway (He et al., 2004; Trauelsen et al., 2021). Likewise, malate is recognized by the succinate receptor, leading to rapid increases in intracellular [Ca^2+^] and IP_3_ accumulation (Trauelsen et al., 2017).

In this study, pharmacological and reverse genetics experiments demonstrated the involvement of second messengers, such as Ca^2+^, cAMP, IP_3_, and G-proteins, in malate signaling in plants (Fig. **7**). Unlike animal GPCRs, plant GPCRs lack well-characterized guanine nucleotide exchange factor activity, and such GPCRs have not yet been identified. Nevertheless, TCA cycle metabolites are common stress-responsive signal molecules mediated by G-protein-dependent signaling cascades in both animal and plant kingdoms.

Based on our findings, we propose a model summarizing malate signaling in guard cells (Fig. **7H**). The malate signal is transmitted *via* G-proteins, which regulate the generation of second messengers. This signaling cascade induces increases in [Ca^2+^]_cyt_, which activates SLAC1 through phosphorylation by Ca^2+^-dependent protein kinases. Consequently, malate promotes Cl^−^ transport through active-form SLAC1, decreasing turgor pressure and driving stomatal closure.

## Materials and Methods

### Plants and growth conditions

Grapevine (*V. vinifera* L. cv. Chardonnay) was grown on a soil mixture of 1:1 = soil: vermiculite (v/v) in a growth room at 24℃ and 80% relative humidity under 16 h-light/8 h-dark photoperiod with a photon flux density of 100 μmol m^−2^ s^−1^. Arabidopsis (*A. thaliana* L. ecotype Colombia-0) was grown on a soil mixture of 1:1 = soil: vermiculite (v/v) in a growth chamber at 21℃ and 60% relative humidity under 16 h-light/8 h-dark photoperiod with a photon flux density of 120 μmol m^−2^ s^−1^. The T-DNA insertion lines, *gpa1* (SALK_001846 and SALK_066823C) and *agb1* (SALK_077086C and SALK_204268C), were obtained from Arashare and NASC.

### Dehydration stress treatment

Fully expanded leaves from 1 to 2-month-old grapevine plants were randomly detached. Dehydration was performed as described previously (Urano et al., 2009) with a few modifications. The detached leaves were exposed to dehydration stress on the paper at 26℃ and ambient humidity under light. At indicated time, the leaves were frozen by liquid nitrogen.

### Sample preparation for LC-MS

The detached leaves were freeze-dried in a lyophilizer (Scientz-100F; Scientz, Zhejiang, China) and then homogenized (30 Hz, 1.5 min) into powder using a grinder (MM 400; Retsch, Dusseldorf, Germany). Next, 1200 μL of -20℃ pre-cooled 70% methanolic aqueous internal standard extract added to 50 mg of sample powder. The sample was vortexed once every 30 min for 30 s, for a total of 6 times. After centrifugation at 12000 rpm for 3 min, the supernatant was aspirated, and the sample was filtered through a microporous membrane (0.22 μm pore size) and stored in the injection vial for UPLC-MS/MS analysis.

### Metabolite analysis using LC-MS

The sample extracts were analyzed using an UPLC-ESI-MS/MS system (UPLC, ExionLC™ AD: SCIEX, MA, USA; MS, Applied Biosystems 4500 Q TRAP: SCIEX). The analytical conditions were as follows, UPLC: column, Agilent SB-C18 (1.8 µm, 2.1 mm * 100 mm); The mobile phase was consisted of solvent A, pure water with 0.1% formic acid, and solvent B, acetonitrile with 0.1% formic acid. Sample measurements were performed with a gradient program that employed the starting conditions of 95% A, 5% B. Within 9 min, a linear gradient to 5% A, 95% B was programmed, and a composition of 5% A, 95% B was kept for 1 min. Subsequently, a composition of 95% A, 5.0% B was adjusted within 1.1 min and kept for 2.9 min. The flow velocity was set as 0.35 mL per min; The column oven was set to 40℃; The injection volume was 4 μL. The effluent was alternatively connected to an ESI-triple quadrupole-linear ion trap (QTRAP)-MS.

The ESI source operation parameters were as follows: source temperature 550℃; ion spray voltage (IS) 5500 V (positive ion mode)/-4500 V (negative ion mode); ion source gas I (GSI), gas II (GSII), curtain gas (CUR) were set at 50, 60, and 25 psi, respectively; the collision-activated dissociation (CAD) was high. QQQ scans were acquired as MRM experiments with collision gas (nitrogen) set to medium. DP (declustering potential) and CE (collision energy) for individual MRM transitions was done with further DP and CE optimization. A specific set of MRM transitions were monitored for each period according to the metabolites eluted within this period.

### Sample preparation for GC-MS

The leaves subjected dehydration stress were ground to a powder in liquid nitrogen. 500 mg (1 mL) of the powder was transferred immediately to a 20 mL head-space vial (Agilent, CA, USA), containing NaCl saturated solution, to inhibit any enzyme reaction. The vials were sealed using crimp-top caps with TFE-silicone headspace septa (Agilent). At the time of SPME analysis, each vial was placed in 60℃ for 5 min, then a 120 µm DVB/CWR/PDMS fiber (Agilent) was exposed to the headspace of the sample for 15 min at 60℃.

### Metabolite analysis using GC-MS

After sampling, desorption of the VOCs from the fiber coating was carried out in the injection port of the GC apparatus (Model 8890; Agilent) at 250℃ for 5 min in the splitless mode. The identification and quantification of VOCs was carried out using an Agilent Model 8890 GC and a 7000D mass spectrometer (Agilent), equipped with a 30 m × 0.25 mm × 0.25 μm DB-5MS (5% phenyl-polymethylsiloxane) capillary column. Helium was used as the carrier gas at a linear velocity of 1.2 mL/min. The injector temperature was kept at 250℃ and the detector at 280℃. The oven temperature was programmed from 40℃ (3.5 min), increasing at 10℃/min to 100℃, at 7℃/min to 180℃, at 25℃/min to 280℃, hold for 5 min. Mass spectra was recorded in electron impact (EI) ionization mode at 70 eV. The quadrupole mass detector, ion source and transfer line temperatures were set, respectively, at 150, 230 and 280℃. The MS was selected ion monitoring (SIM) mode was used for the identification and quantification of analytes.

### Data analysis of the non-target metabolome

Relative metabolite abundances were calculated by the peak areas. Unsupervised PCA was performed by statistics function prcomp within R (www.r-project.org). The relative contents of all differential metabolites were processed by UV (unit variance scaling) followed by K-Means cluster analysis. Identified metabolites were annotated using KEGG Compound database (http://www.kegg.jp/kegg/compound/). Pathways with significantly regulated metabolites mapped to were then fed into metabolite sets enrichment analysis (MSEA), their significance was determined by hypergeometric test’s P-values. Upregulated metabolites are defined VIP >1, log_2_ (Fold change) ≥1 and P-value >0.05, downregulated metabolites are defined VIP >1, log_2_ (Fold change) ≤−1 and P-value >0.05. P-value was calculated by Welch’s t-test.

### [Ca^2+^]_cyt_ imaging

Wild-type Arabidopsis plants expressing Yellow Cameleon 3.6 were used to measure [Ca^2+^]_cyt_ in guard cells as described previously (Mimata et al., 2022b). The abaxial side of an excised rosette leaf was gently attached to a glass slide with a medical adhesive (stock no. 7730; Hollister, IL, USA) and then mesophyll tissues were whittled away with a razor blade to keep the abaxial epidermis intact on the slide. The remaining abaxial epidermis was immersed in stomatal assay solution, comprising 5 mM KCl, 50 μM CaCl_2_ and 10 mM MES/Tris (pH 5.6), in the light for 2 h to induce stomatal opening. The epidermis was treated with 10 mM TCA cycle metabolites in stomatal assay solution at the indicated time. Inhibitors were added 5 min before starting imaging. The stock solution of TCA cycle metabolites was dissolved in stomatal assay solution and adjusted to a pH of 5.6 with Tris.

The images were acquired under a fluorescence microscope (ECLIPSE Ti2-E; NIKON). Excitation light was provided by a mercury arc lamp and a 436 nm filter (ET436/20x, Chroma Technology Corporation, VT, USA). Emission of the CFP was measured at 480 nm filter (ET480/40m, Chroma Technology Corporation) and of the YFP at 535 nm filter (ET535/30m, Chroma Technology Corporation) using a CMOS camera (ORCA-Fusion BT Digital CMOS camera C15440; HAMAMATSU, Shizuoka, Japan). Images were taken every 5 s.

### Modeling

AlphaFold3 was used to predict the protein structure of VvSLAC1. Five protomer models were generated, with predicted template modeling scores raging from 0.65 to 0.67 and ranking scores raging from 0.81 to 0.83. The models were compared with the cryo-EM structure of AtSLAC1 (PDBs 8gw6: Lee et al., 2023) and the top-scored model has a root mean square deviation of 0.852 Å/335 Cα. Due to the limited length of the AtSLAC1 structure, the residues 1-141 and 507-553 in VvSLAC1 were removed.

### Cloning and cRNA synthesis

All constructs were cloned into the oocyte expression vector pNB1u (Nour-Eldin et al., 2006) by ClonExpress II One Step Cloning Kit (Vazyme Biotech, Nanjing, China). The site-directed mutants were generated by FastCloning (Li et al., 2011). VvSLAC1 (LOC100244459) cDNA from *V. vinifera* was used for cloning, and all constructs were verified by sequencing. Primers used for cloning and site-directed mutagenesis are listed in Table S3. cRNA was prepared using an mMESSAGE mMACHINE ^TM^ T7 Transcription Kit (Thermo Fisher Scientific, MA, USA).

### Two-electrode voltage-clamp

*Xenopus laevis* oocytes were injected with 50 nL cRNA (each 10 ng) and incubated in ND96 buffer at 18℃ for a few days before voltage-clamp recordings (Mimata et al., 2022b). The bath solution contained 1 mM Mg-gluconate, 1 mM Ca-gluconate and 1 mM LaCl_3_ ± 10 mM TCA cycle metabolites buffered with 10 mM MES/Tris to adjust the pH to 5.6. Osmolality was adjusted to 220 mOsmol kg^−1^ using D-sorbitol. The voltage pulse was commanded to clamp the membrane potential at −120 mV in gap-free or from +60 to −160 mV in 20 mV decrements in step for 2.5 s with a holding potential of 0 mV. Voltage-clamp recordings for oocytes were performed using an Axoclamp 900A amplifier (Molecular Devices, CA, USA), data were acquired using a Digidata 1550B system (Molecular Devices) and analyzed using pCLAMP 11.2 software (Molecular Devices).

### Measurement of stomatal aperture

Stomatal apertures were measured as described previously (Ye et al., 2020) with modifications. Leaf discs (4 mm in diameter) obtained from fully expanded leaves were placed abaxial side down on stomatal assay solution. The discs were exposed to light for 2 h to induce stomatal opening and subsequently treated with 10 mM TCA cycle metabolites in stomatal assay solution for an additional 2 h. Inhibitors were added 5 min before malate treatment. The types and concentrations of inhibitors are listed in Supplementary Table S2. The abaxial epidermis was captured under optical microscopes (ECLIPSE Ts-2R, ECLIPSE Ti2-E and ECLIPSE Ci; NIKON, Tokyo, Japan) using NIS ELEMENTS software (NIKON). Stomatal apertures were quantified using IMAGEJ software (NIH). We measured 30 stomatal apertures from a leaf disc to calculate an average. This measurement was repeated four times using different plants, and the overall average was calculated.

### Measurement of ROS production

ROS production in guard cells was analyzed using the fluorescent dye 2’,7’-dihydrodichlorofluorescein diacetate (H_2_DCF-DA) as described previously (Mimata et al., 2022a) with modifications. The abaxial side of an excised rosette leaf was gently attached to a glass slide with a medical adhesive and then mesophyll tissues were whittled away with a razor blade. The remaining abaxial epidermis was immersed in stomatal assay solution in the light for 2 h to induce stomatal opening. A total of 50 μM H_2_DCF-DA was added to the stomatal assay solution and the epidermal tissues were incubated in the dark for 30 min. After the dye loading, the epidermal tissues were gently rinsed with stomatal assay solution. The epidermis was treated with 10 mM TCA cycle metabolites ± inhibitor in stomatal assay solution. After the 30 min incubation, fluorescent signals were captured using the fluorescence microscope with 480 ± 15 nm/ 535 ± 23 nm excitation/emission filters. We measured 30 guard cells from an epidermis to calculate an average. This measurement was repeated three times using different plants, and the overall average was calculated.

## Acknowledgements

We thank Dr. Hussam Hassan Nour-Eldin (University of Copenhagen) for providing pNB1u vector; Dr. Yoshiyuki Murata and Dr. Shintaro Munemasa (Okamaya University) for providing plasmid constructs; Dongyue Wang (Peking University Institute of Advanced Agricultural Sciences) for helping cloning.

## Authors’ contributions

YM and WY planned and designed the research. YM, RG and XP performed experiments. YM analyzed data. YM, GQ and WY interpreted the data. YM and WY wrote the manuscript.

## Ethics approval and consent to participate

Not applicable.

## Consent for publication

All authors approve the manuscript and consent to the publication of the work.

## Availability of data and materials

The datasets during and/or analyzed during the current study available from the corresponding author on reasonable request.

## Supplementary information

The following materials are available in the online version of this article.

**Supplementary Figure S1.** Metabolome analysis in grapevine leaves during dehydration treatment.

**Supplementary Figure S2.** Ca^2+^ response to acetate in guard cells.

**Supplementary Figure S3.** Malate-induced stomatal closure in the presence of inhibitors.

**Supplementary Figure S4.** ROS production in the presence of TCA cycle metabolites.

**Supplementary Table S1.** Dataset of metabolome analysis.

**Supplementary Table S2.** List of inhibitors used in this work.

**Supplementary Table S3.** List of primers used in this work.

