## Supplemental Figures for "A drought stress-responsive metabolite malate modulates stomatal responses through G-protein-dependent pathway in grapevine and Arabidopsis"

This file includes Supplementary Figure S1–4 and Supplementary Table S2–3.

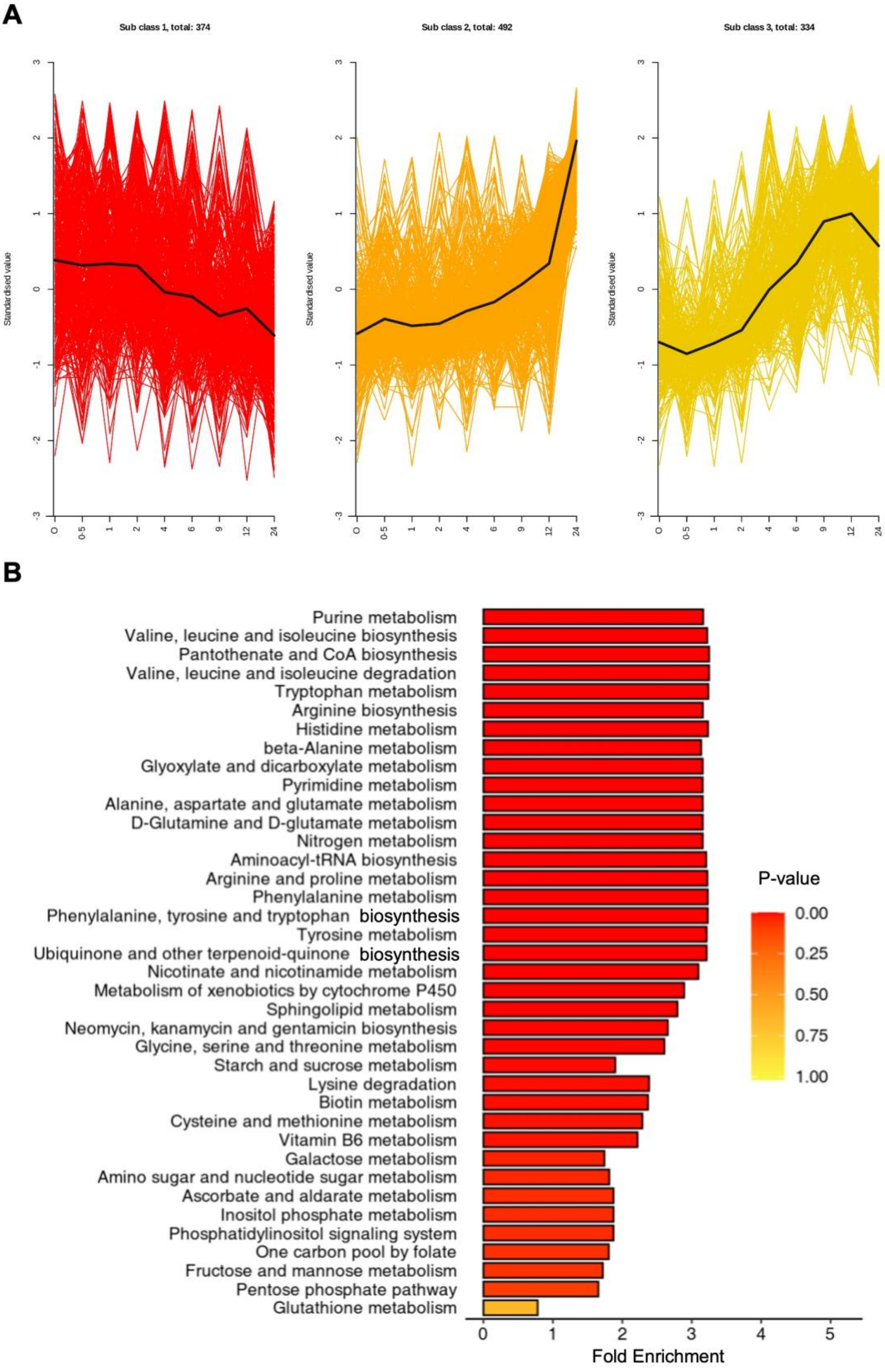

A
Sub class 1, total: 374
Sub class 2, total: 492
Sub class 3, total: 334
Standardised value
0
0-5
1
2
4
6
9
12
24
B
Purine metabolism
Valine, leucine and isoleucine biosynthesis
Pantothenate and CoA biosynthesis
Valine, leucine and isoleucine degradation
Tryptophan metabolism
Arginine biosynthesis
Histidine metabolism
beta-Alanine metabolism
Glyoxylate and dicarboxylate metabolism
Pyrimidine metabolism
Alanine, aspartate and glutamate metabolism
D-Glutamine and D-glutamate metabolism
Nitrogen metabolism
Aminoacyl-tRNA biosynthesis
Arginine and proline metabolism
Phenylalanine metabolism
Phenylalanine, tyrosine and tryptophan biosynthesis
Tyrosine metabolism
Ubiquinone and other terpenoid-quinone biosynthesis
Nicotinate and nicotinamide metabolism
Metabolism of xenobiotics by cytochrome P450
Sphingolipid metabolism
Neomycin, kanamycin and gentamicin biosynthesis
Glycine, serine and threonine metabolism
Starch and sucrose metabolism
Lysine degradation
Biotin metabolism
Cysteine and methionine metabolism
Vitamin B6 metabolism
Galactose metabolism
Amino sugar and nucleotide sugar metabolism
Ascorbate and aldarate metabolism
Inositol phosphate metabolism
Phosphatidylinositol signaling system
One carbon pool by folate
Fructose and mannose metabolism
Pentose phosphate pathway
Glutathione metabolism
0 1 2 3 4 5
Fold Enrichment
P-value
0.00
0.25
0.50
0.75
1.00

**Supplementary Figure S1. Metabolome analysis in grapevine leaves during dehydration treatment.**

**A)** K-Means plot of differential metabolite. Black lines indicate the means in each group. The ordinate represents the standardized relative content of metabolites. Sub class represents the metabolite category number with the same changing trend, with total: # indicating the number of metabolites in this category.

**B)** MSEA of differential metabolites. Bar color indicates the significance of difference, and bar lengths represent the fold enrichment. Significance was determined by hypergeometric test's P-values.

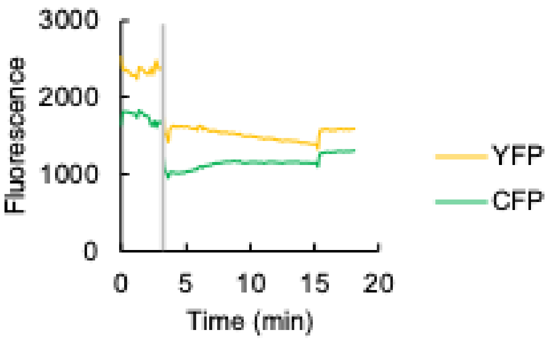


**Supplementary Figure S2. $Ca^{2+}$ response to acetate in guard cells.**

Representative traces of fluorescence emission intensity in *A. thaliana* guard cells expressing the $Ca^{2+}$ sensor Yellow Cameleon 3.6. Grey bars indicate the time point when treatment was applied.

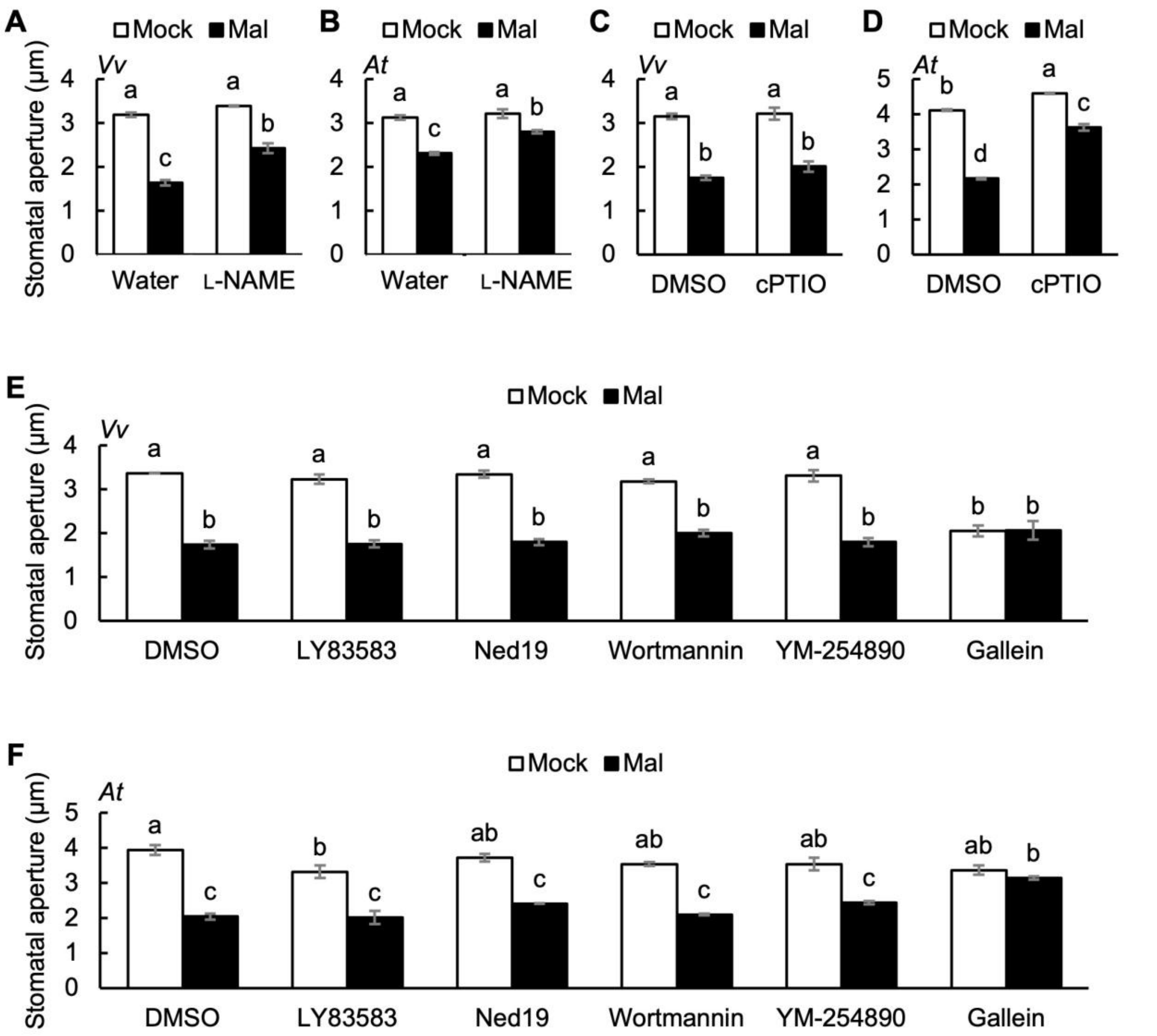


**Supplementary Figure S3. Malate-induced stomatal closure in the presence of inhibitors.**

**A to F)** Effects of inhibitors on malate-induced stomatal closure in **A, C and E)** *V. vinifera* or **B, D and F)** *A. thaliana* leaves. Averages of stomatal apertures from four independent experiments ($n = 4$) are shown. Data are the mean ± SE. Different letters indicate statistical significances based on two-way ANOVA with Tukey's HSD test, $P < 0.05$.

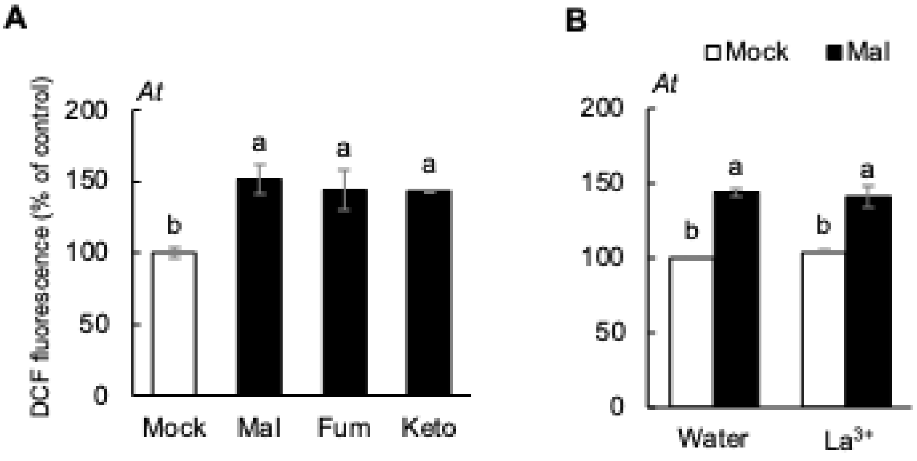


**Supplementary Figure S4. ROS production in the presence of TCA cycle metabolites.**

**A)** Effects of TCA cycle metabolites on ROS production in *A. thaliana* guard cells.

**B)** Effects of $La^{3+}$ on malate-induced ROS production in *A. thaliana* guard cells. The ROS-sensitive dye, 2',7'-dichlorodihydrofluorescein diacetate ($H_2$DCF-DA) was used for ROS detection in guard cells. Fluorescence intensity was normalized to mock value **A)** or mock value in water **B)**. Averages from three independent experiments (n = 3) are shown. Data are the mean ± SE. Different letters indicate statistical significances based on one-way ANOVA with Tukey's HSD test **A)** or two-way ANOVA with Tukey's HSD test **B)**, $P < 0.05$.

**Supplementary Table S2. List of inhibitors used in this work.**

| **Name** | **Inhibitor type** | **Final concentration** |
|---|---|---|
| DIDS | Anion channel blocker | 100 μM |
| 9-AC | Anion channel blocker | 100 μM |
| BAPTA | Extracellular $Ca^{2+}$ chelator | 100 μM |
| Nifedipine | $Ca^{2+}$ channel blocker | 10 μM |
| $La^{3+}$ ($LaCl_3$) | $Ca^{2+}$ channel blocker | 1 mM |
| Nicotinamide | cADPR synthesis inhibitor | 50 mM |
| Alloxan | cAMP synthesis inhibitor | 1 mM |
| Neomycin | $IP_3$ synthesis inhibitor | 50 μM |
| SHAM | Peroxidase-catalyzed ROS production inhibitor | 2 mM |
| L-NAME | NO synthetase inhibitor | 25 μM |
| cPTIO | NO scavenger | 100 μM |
| LY83583 | NO-sensitive guanylate cyclase inhibitor | 2 μM |
| Ned 19 | NAADP antagonist | 10 μM |
| Wortmannin | $PIP_3$ synthesis inhibitor | 5 μM |
| YM-254890 | G-protein inhibitor | 10 μM |
| Gallein | G-protein inhibitor | 10 μM |
| GDPβS | G-protein inhibitor | 10 μM |
| GRK2i | G-protein inhibitor | 5 μM |
| Suramin | G-protein inhibitor | 100 μM |

**Supplementary Table S3. List of primers used in this work.**

| Name | Sequence |
|---|---|
| VvSLAC1_pNB1u_F | aaacctcagcgaattcATGGACAGAAGACCGACTTC |
| VvSLAC1_pNB1u_R | gcctattcccaagcttTCAGTGCTCTGCTTCCTTC |
| VvSLAC1F440A_F | TACACCgctCCCATGACAACAGTATCAGTGGC |
| VvSLAC1F440A_R | CATGGGagcGGTGTAAGACCACCATGCCAC |
